# A layered standards framework for integrating single-cell and spatial omics data into brain cell atlases

**DOI:** 10.64898/2026.04.30.722039

**Authors:** Patrick L. Ray, Jeremy A. Miller, Dorota Jarecka, Kimberly A. Smith, Pamela M. Baker, Lydia Ng, Maryann E. Martone, Puja Trivedi, Rashmie Abeysinghe, Lisa Anderson, Anita E. Bandrowski, Edyta Vieth, Ashwin A. Bhandiwad, Tek Raj Chhetri, Licong Cui, Michelle Giglio, Jeff Goldy, Na Hong, Hao Huang, Yan Huang, Yasmeen Hussain, Nelson J. Johansen, Mariah Kenney, Lauren Kruse, Xiaojin Li, James C. Meldrim, Tyler Mollenkopf, Suvarna Nadendla, David Osumi-Sutherland, Raymond Sanchez, Richard H. Scheuermann, Shiqiang Tao, Charles R. Vanderburg, Yuntao Yang, Alex Ropelewski, Shoaib Mufti, Ed S. Lein, Hua Xu, W. Jim Zheng, Satrajit S. Ghosh, Owen White, Michael Hawrylycz, Guo-Qiang Zhang, Carol Thompson

## Abstract

The BRAIN Initiative Cell Atlas Network (BICAN) is generating large-scale multimodal datasets to profile cell types in the human, non-human primate, and mouse brain. The diversity of single-cell and spatial transcriptomic and epigenomic assays, combined with varied experimental contexts, multiple data-generating laboratories and distributed infrastructure, poses substantial challenges for data integration and reuse in BICAN. To address this, we implemented a standards framework that enables layered integration of these data into knowledge-ready products for interoperable brain cell atlases. This framework organizes data based on three progressively structured layers. First, we introduced an assay-agnostic modeling layer that unifies the representation of single-cell and spatial omics data using a common set of biological entities and processes assessed by diverse experimental techniques. Second, we implemented harmonized metadata standards that capture key experimental features linked to biospecimen provenance across heterogeneous tissue sources, species, and preparations, supporting integration and validation while minimizing burden on data contributors. Third, we present an extensible representation for data-driven cell type taxonomies that integrates molecular data with annotations, ontology mappings, and evidence. Together, these contributions represent an end-to-end framework that transforms heterogeneous datasets into structured, interoperable resources that support broad community reuse via mapping algorithms, annotation systems, and visualization platforms. This approach links biospecimen provenance with cell-level outputs and embeds these in a standardized taxonomy format, enabling downstream applications such as cross-dataset integration, reference mapping, and knowledge-driven analysis. More broadly, our work demonstrates a generalizable strategy for enabling an efficient data-to-knowledge pipeline in a large-scale consortium setting.

## 1. Introduction

In brain cell atlasing, rapid advances in single-cell and spatial molecular profiling technologies have expanded the scale, multimodality, and resolution at which cellular diversity can be measured. These technologies produce high-dimensional datasets that now serve as the basis for cell type discovery and classification. As efforts such as the BRAIN Initiative Cell Atlas Network (BICAN; brain-bican.org) generate cell typing data for hundreds of millions of cells across species and developmental stages, with 17 million cells profiled by sequencing-based methodologies in the basal ganglia alone (BRAIN Initiative Cell Atlas Network et al., 2026), the integration of these data with cell type knowledge requires consistent representation of experimental provenance, biospecimen metadata and knowledge structures across scientific and infrastructure partners.

Genome-scale projects have demonstrated the power of standardization across large collaborative efforts to achieve historic advances. The success of initiatives such as the Human Genome Project(Lander et al. 2001) and the ENCODE Project (Dunham et al. 2012) relied on shared data models, coordinated metadata practices and interoperable infrastructures essential for large-scale integrative analysis. In neuroscience, efforts to map cellular diversity in human, non-human primate and mouse brains have entered a similarly data-intensive phase, driven by single-cell and spatial omics technologies. The need for coordinated standards has begun to yield alignment on neuroanatomy, computational pipelines, and coordinate systems(Abrams et al. 2022). Current community roadmaps for single-cell atlas efforts also highlight the importance of metadata for the integration of single-cell atlases(Haniffa et al. 2021; Zilbauer et al. 2023). However, the rapidity at which emerging technologies continue to add depth, complexity, multimodality, and scale requires adaptable strategies for integrating new diverse data to support a growing reference atlas of cell types in the brain.

A broad community effort to develop a cell census of the brain faces three core standardization challenges: heterogeneity across assay types and measurement modalities, inconsistent metadata arising from diverse species, protocols, tissue sources and laboratories, and inconsistent representations of biological knowledge such as cell type taxonomies, annotations and evidence. Standards for gene expression and single-cell experiments have evolved for more than two decades to address aspects of these challenges. For example, the Minimum Information About a Microarray Experiment (MIAME) standard(Brazma et al. 2001) helped establish the value of standardized experimental reporting for interpretation, reanalysis and reuse of gene expression data. More recently, the Minimum Information about a Single-Cell Experiment (minSCe) guidelines have been proposed for robust comparative analyses of scRNA-seq(Füllgrabe et al. 2020). These standards are tailored to individual assay types and support a broad swath of reanalysis use cases, but do not fully address cross-assay metadata and knowledge-representation needed for an integrated brain cell atlas.

While minimal information standards are valuable for establishing a baseline level of metadata consistency across experiments, they may be insufficient on their own for capturing detailed provenance, biospecimen relationships and protocol information that may be critical for interpretation and reuse. The Ontology of Biomedical Investigations (OBI)(Brinkman et al. 2010; Vita et al. 2021) provides a complementary framework for modeling biomedical investigations, including experimental entities, processes, inputs and outputs. This abstraction supports flexible representation across diverse biomedical experiment types and well suited for expert-driven ontology development. However, because OBI is intentionally broad and non-prescriptive, additional community agreement is needed to define how its concepts should be instantiated for brain cell atlas data. A shared, purpose-driven metadata instantiation can unify information across techniques and laboratories while providing sufficient structure to support common workflows, validation, downstream analysis and visualization and cross-modality integration.

In parallel, data formats for representing single-cell measurements have evolved to support large-scale analysis and sharing. Matrix-based schemas, such as the AnnData format(Virshup et al. 2024), have been developed to represent cell-by-feature data and metadata and support interoperability across tools. Community efforts have further provided standards for cell- and gene-level annotation(Chan Zuckerberg Initiative [2020] 2026) for data sharing, but still lack formal structure for taxonomy versioning, inclusion of novel cell types, and explicit tracking of evidence. With recent approaches to define cell types directly from large-scale data, the need for standardized mechanisms to cross-reference cells and data to novel provisional classifications and layers of knowledge becomes increasingly important.

BICAN, a consortium with over 700 participating researchers, is funded by the NIH BRAIN Initiative to create reference brain atlases cataloging cell types across species, brain regions, and developmental stages using transcriptomics, epigenomics, proteomics, morphoelectric characterization, and spatial profiling(Network (BICAN) et al. 2026). BICAN comprises multiple data generating labs, sequencing cores, data repositories and data pipelines, integrated into a federated ecosystem (**Fig. S1**). Large-scale consortium efforts require a unified framework that links data, provenance, and ultimately knowledge to create integrated cell atlases and enable broad re-use of the resulting comprehensive cell type atlases by researchers and artificial intelligence (AI) systems alike.

Here we describe the multi-layered approach to standardizing metadata and knowledge representation in BICAN. First, we introduce an assay-agnostic modeling layer that unifies the representation of single-cell and spatial omics data using a common set of biological entities, or biospecimens, and processes to describe diverse experimental techniques. This model serves as a backbone for the development of detailed metadata for sequencing library generation steps from tissue to sequence, suitable for capturing entity provenance in a common way across these methods. Second, we implement harmonized metadata standards that capture experimental conditions across heterogeneous tissue sources, species, and preparations, supporting integration and validation while minimizing burden on data contributors. This solution maximizes accommodation for heterogeneous tissue inputs with minimal burden, using a pragmatic, inclusive schema design. Third, we present an extensible representation for data-driven cell type taxonomies that integrates molecular data with annotations, ontology mappings, and evidence. This cell type taxonomy and annotation standard elevates the matrix-based CELLxGENE data model to integrate derived biological knowledge. This data structure provides a novel solution to merging data with knowledge into a common framework to support cross-analysis between a cell type reference and datasets generated by other users.

## 2. Design

### Governance and Workflow

BICAN operations are managed within a governance framework that utilizes topic-specific working groups (WGs) that coordinate cross-lab and cross-institution activities. Standards development begins with subject matter experts working in concert with the BICAN’s Metadata and Ontologies Working Group and Data Ecosystem Working Group to co-develop, integrate, and document standards for review and approval by the consortium’s BOSC Operations Steering Committee (see **Fig. 1A**). Implementation of donor and biospecimen metadata standards occurs through the Neuroanatomy-anchored Information Management Platform for Collaborative BICAN Data Generation portal (NIMP; RRID:SCR_024684)(Tao et al. 2025), which manages tissue requests, submission of sequencing libraries, and deposition of sequencing data with QC. Standards-compliant metadata then flow through to final data release through modality-specific data archives. Dissemination of data products adhering to standards represents an integral feature of BICAN data.

**Figure 1.**
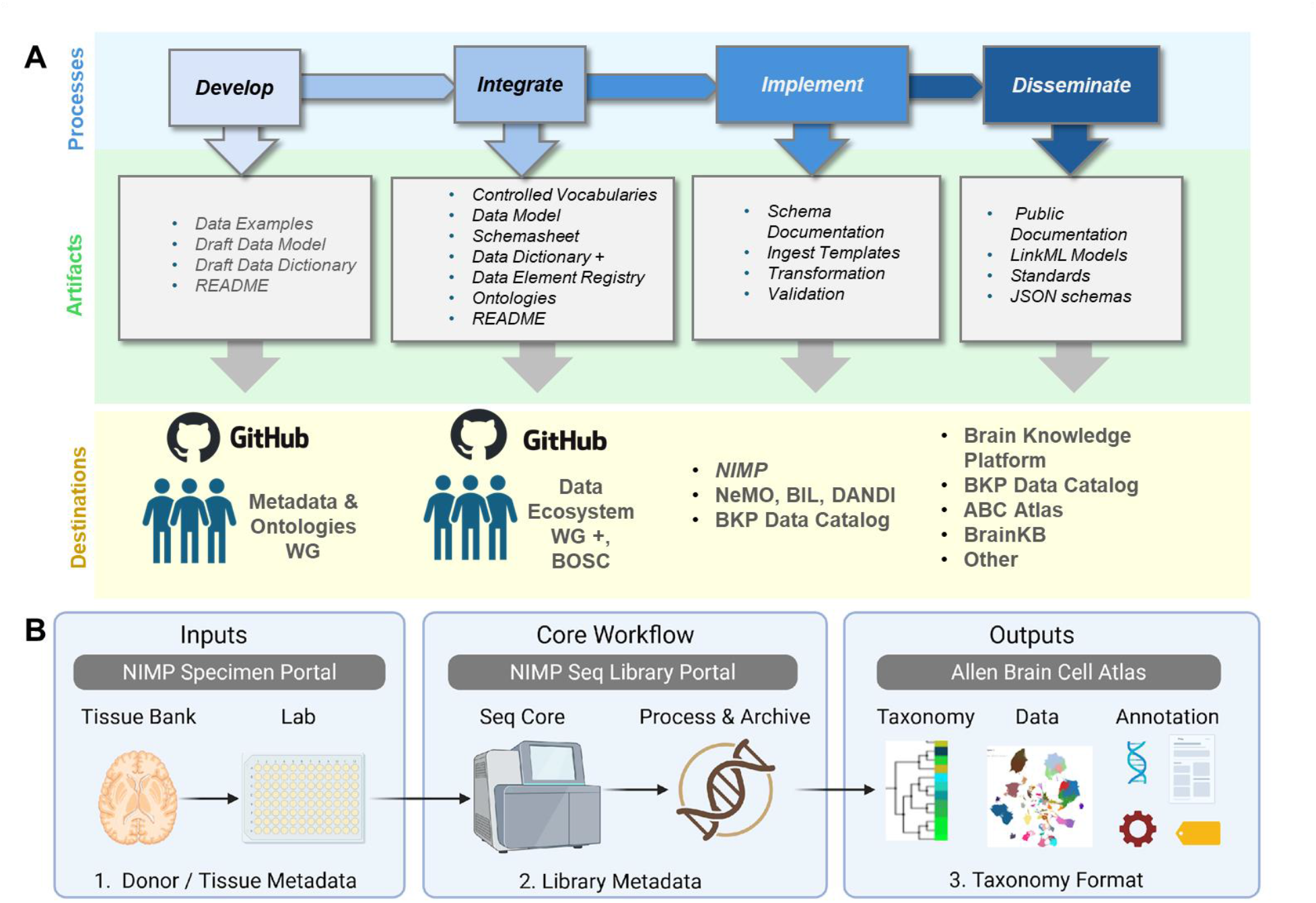
Overview of the development pipeline for BICAN data standards and their integration into core BICAN infrastructure. A) The iterative process for developing, integrating, implementing, and disseminating metadata and taxonomy standards. During development, the Metadata and Ontologies WG collaborates with subject matter groups to develop and refine schemas and standards. Documentation artifacts for integrated standards are provided for review and approval by the Data Ecosystem WG for cross-archive data management and integration, and by the BICAN Operations Steering Committee (BOSC) to be considered for official endorsement within the consortium. Artifacts at each stage are deposited into an appropriate GitHub repository for upstream steps and integrated into public resources for downstream steps. B) Illustration of how three metadata standards cover the flow of tissue to data to knowledge. (1) Donor and tissue metadata enter the system from tissue banks and laboratories for experiments; (2) Library metadata generated, processed, and stored in centralized workflows; and (3) the taxonomy format used to represent data-driven cell type taxonomies within the Allen Brain Cell Atlas and related knowledge platforms.

Metadata schema development required collaboration between scientific subject matter experts (SMEs) and engineering representatives to enable data representation with scientific fidelity in robust systems (**Fig. 1A**). At a high level, the process consisted of an inventory of the breadth of metadata fields and parameters through iterative collaboration between those SMEs and the engineers; a period of metadata integration, including assessment of mapping to known NIH Common Data Elements (Wang et al. 2025), and mapping across the distributed BICAN data ecosystem. The review and approval of the schema standard utilized the BICAN governance framework; however, continuing updates included addition, removal, and edits of metadata fields over time that were implemented in NIMP and recorded in the schema repository. This approach resulted in the selection of metadata that was both useful for scientific analysis and NIMP tracking. Other infrastructure components disseminate data to the community, including the Neuroscience Multiomic Archive (NeMO Archive, RRID:SCR_016152) (Ament et al. 2023), the Brain Image Library (BIL, RRID:SCR_017272) (Kenney et al. 2024) and the Distributed Archives for Neurophysiology Data Integration (DANDI, RRID:SCR_017571). Datasets are listed in the BICAN Data Catalog (RRID : SCR_027884) accessed via the Allen Institute Brain Knowledge Platform (RRID:SCR_027180).

### Input Artifacts and Curation

Relevant metadata fields were initially provided by scientists as a spreadsheet or other text-based tabular documentation. The metadata definitions were converted into a standard Data Dictionary format (Sharma et al. 2016) that formally documented metadata elements with essential usage information, including definitions, data types, controlled vocabulary requirements, and alternative terms or labels. Controlled vocabularies were documented and terms were harmonized across partner data repositories and standard ontologies. After an iterative process of improvement, the documentation was added to the public GitHub repository (https://github.com/brain-bican/metadata-schemas) and reviewed by the BICAN community. All approved models were versioned and made available in the repository for use by the broader community.

### Entity-centric data models for molecular profiling

Data models were created to represent biological specimens as key entities unified across both single-cell and spatial molecular profiling. These models were collaboratively and iteratively generated within the working group, and the approach is conceptually aligned with object-oriented domain modeling, in which stable entities and their relationships form the backbone of a system representation. As part of the curation process, specimen and process entities from the data models were linked to their associated metadata fields in the schema data dictionary.

### Formal, Machine-Readable Data Models

Metadata fields and controlled vocabularies from the metadata-schemas GitHub repository were converted into formal, machine-readable data models using LinkML tools (Moxon et al. 2021), and shared in the public GitHub repository: https://github.com/brain-bican/models. Provenance components were added to the LinkML schemas to align with the entity-centric, biospecimen provenance data models. These formal schemas serve several critical functions: (i) validation of incoming data, (ii) transformation of heterogeneous data into a consistent representation, and (iii) enabling interoperability of data across downstream tools and platforms. By embedding fields for persistent identifiers and provenance information directly into the schema, and by supporting links to external ontologies (Unni et al. 2022) where appropriate, the schemas provide a structured framework for recording dataset provenance, and representing relationships between data and derivatives. LinkML’s capacity to represent complex, interlinked schemas makes it particularly well suited for large-scale, evolving metadata standards.

To operationalize these schemas, we developed *bkbit* (RRID:SCR_027564), a Python library of data translators that convert domain-specific data into standardized, schema-compliant JSON-LD(Kellogg et al. 2019). Each translator was tailored to a specific schema and ensures conformity with consortium-wide metadata standards. The resulting JSON-LD files embed semantic meaning, are human-and machine-readable, and support downstream ingestion into diverse services. Contributors submitted raw data and bkbit handles parsing, transformation, and validation in a reproducible and scalable manner.

### Outputs, dissemination, and availability

We developed and finalized each metadata standard to support biological data integration and interoperability in BICAN. These standards addressed critical domains spanning anatomy, cellular annotation and taxonomy formats, metadata for donor information and experimental details, among others (**Table S1)**. Each metadata standard was accompanied by supporting documentation designed to facilitate implementation and ensure consistent usage across research groups. Standards are maintained in a public version-controlled GitHub repository (https://github.com/brain-bican/metadata-schemas) providing full transparency of development history, issue tracking, and community contribution mechanisms. Each release is tagged with semantic versioning to enable reproducible implementations. The NIMP terminology browser (https://brain-specimenportal.org/terminology_browser ; account required) allows investigators to interactively explore and inspect the BICAN data elements and controlled terms in use in the NIMP portal. All the LinkML schemas are also shared using GitHub: https://github.com/brain-bican/models.

## 3. Assay-agnostic modeling to support biospecimen provenance, single-cell and spatial assay metadata

A core feature of the BICAN consortium is a common single-cell and spatial sequencing workflow that uses shared molecular pipelines for reference genome alignment, quality metric generation, and the production of standardized data artifacts, despite data originating from multiple laboratories and diverse assay types. These assays include single-cell transcriptomic, chromatin accessibility, and methylation, and multimodal methods such as 10x Multiome (RNA and chromatin accessibility) and Patch-seq (RNA, morphological reconstruction, and patch-clamp electrophysiology). Furthermore, imaging-based and sequencing-based spatial transcriptomics methods have emerged that provide anatomic context with single-cell or near-single-cell resolution of gene expression. The diversity and complexity of methodologies (subset listed in **Table S2**) presented challenges for standardization. Because both primary processing of sequencing files and downstream analysis relies on upstream metadata about sample preparation, BICAN required more systematic data organization of sample, library, and provenance metadata. Therefore, we developed a unified framework that is generalizable for most single-cell sequencing techniques and is extensible to related sequencing-based spatial assays.

BICAN centralized most single-cell sequencing submission through a shared workflow, in which individual laboratories and a designated sequencing cores utilize the NIMP portal to manage submissions, metadata, and quality control steps. This made it essential to define, control, and standardize the sample, library, process, and QC metadata across the consortium. To support metadata harmonization in a central metadata repository, a task force was convened to collect key requirements from stakeholders, including representatives of data generating laboratories, sequencing cores, and software engineers. The first step for the task force in metadata harmonization included the collection of key fields and their possible values from the relevant stakeholders. These requirements ensured that the resulting schema could accommodate diverse assays while capturing information important for downstream processing and data pipelines.

Because the diversity of experimental assays made it challenging to align experimental steps across laboratories, the next step was to explicitly model experimental processes to provide a framework for understanding the requirements. During the design of the library metadata schema, the engineers used Unified Modeling Language (Shegogue and Zheng 2005) to formally represent, analyze, and capture key components of the library generation process in detail. In parallel, domain experts (neuroscientists) developed a visual process diagram that carefully mapped out the physical form of a sample as it progressed through the workflow accompanied by experimental protocols that produced them so that everyone understood the steps involved. Throughout this collaborative effort, informaticians and domain experts exchanged information continuously, iteratively refining both models. These parallel modeling approaches evolved in tandem and converged as the Library Metadata specification was finalized, ensuring alignment between technical structure and biological context.

A key output of this process was a set of abstract, high level models representing different classes of assays that were generalizable to all single-cell sequencing assays, and conceptually consistent with the experimental process abstraction of OBI (Brinkman et al. 2010). The resultant model allowed researchers and engineers to identify the primary entities (i.e., biospecimens) in the workflow and the processes that derive them, and enabled identification of key entities to which properties and resultant data files are assigned. This model was later extended to also represent spatial transcriptomics experiments, which assign gene expression to anatomic context in the tissue through either a sequencing- or imaging-based approach, and for patch-seq experiments, which utilized a combination of whole cell patch electrophysiology, morphology, and single nucleus sequencing. Altogether the models are generic enough to support a wide range of different single-cell techniques (**Table S2)**.

At an abstract level, single-cell sequencing experiments can be represented as a series of transformations of biological entities connected by defined laboratory processes (**Fig. 2**). A biospecimen, such as a piece of fresh or frozen tissue, undergoes processing which may include some combination of dissection, sectioning, dissociation or nuclei isolation. Cells or nuclei are then captured, isolated or enriched through methods such as flow cytometry, FACS sorting, microfluidics, or other techniques to create a dissociated cell sample. Assay-specific chemistry is applied to incorporate molecular barcodes (i.e., “barcoded cell sample”) to uniquely identify cells or samples and associate molecular measurements with individual cells, nuclei, samples or spatial capture locations. These steps produce assay-specific sequencing libraries which may be divided into library aliquots, pooled for sequencing, and processed at sequencing cores. Downstream computational pipelines produce files that associate the readout (gene counts, methylation status, chromatin accessibility) per cell.

**Figure 2.**
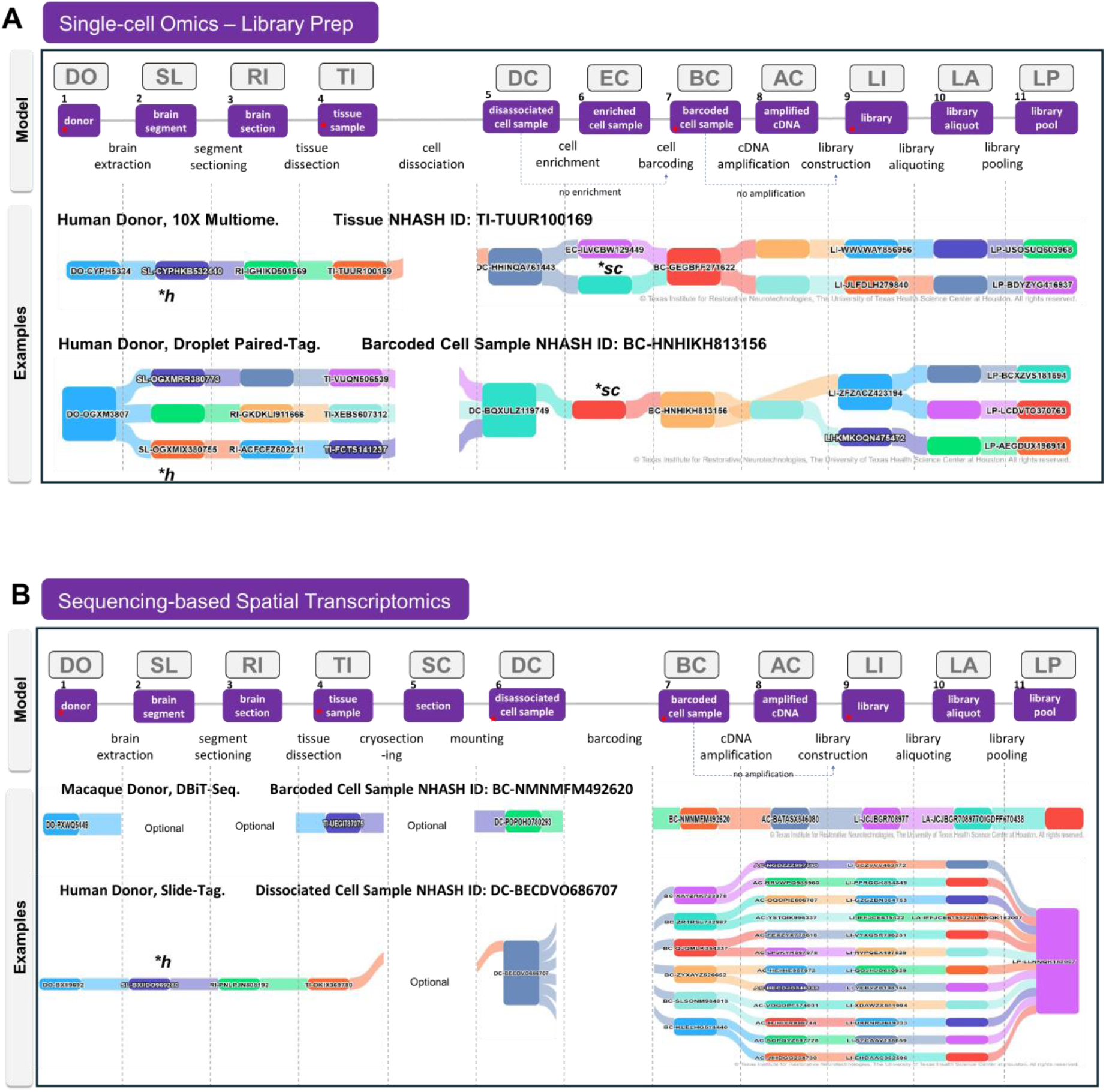
Abstract biospecimen provenance model to support maximal integration across diverse assays. **A**, Model for biospecimen processing generalized for all single cell omics assays. Biospecimen entities are shown in purple boxes with the process steps interleaved. Downstream sequencing and analysis steps not shown. Above each biospecimen, the NHASH identifier abbreviation is shown: DO, donor; SL, slab or brain segment; RI, Region of interest or brain segment section; TI, tissue sample; SC, tissue section or cryosection; DC, dissociated cell sample; EC, enriched cell sample; BC, barcoded cell sample; AC, amplified cDNA; LI, library; LA, library aliquot; and LP, Library Pool. Two examples of biospecimen provenance graphs are shown utilizing visualizations from the NIMP Resource Browser, for 10X Multiome experiments and for Droplet Paired-Tag. The NHASH ID for the anchor entity used for the visualization graph is listed. **B**, Sequencing-based spatial transcriptomics is aligned with the single-cell omics model. The major difference is the introduction of a cryosection biospecimen, and the omission of an enriched cell sample. Examples of biospecimen provenance are shown here for DBiT-Seq and Slide-Tag. Red stars indicate conservation of entities across all assays in that category. Each column of the graph represents one of eleven levels of specimen available for visualization, starting with Donor (DO) and ending with Library Pool (LP). Breaks are shown in the river plots to highlight the absence of certain specimen types. Slabs are typically associated with human samples (*h), and enriched cell samples are optionally available for assays with dissociated single cells or nuclei (*sc). See supplemental Table S1 for a more detailed description of how this model fits multiple assay types.

Furthermore, we were able to align sequencing-based spatial transcriptomics methods, such as Slide-Tag or DBiTseq, with the same basic model by adding a “section” entity that is placed upon a slide or coverslip “substrate”. In the provenance model, the substrate occupies a position analagous to the “dissociated cell sample”, while spatially indexed assay chemistry generates the equivalent of the “barcoded cell sample” by associating molecularly barcodes with spatial location in the tissue. Similarly, imaging-based spatial transcriptomics (e.g., MERFISH), utilizes a “section” entity that is placed on a slide substrate, followed by in situ hybridization and sequential imaging with a gene probe panels. Although these methods do not produce barcoded cell samples in the same way as sequencing-based assays, they can be represented in the model as analagous entities. While the original model was an entity-process diagram, the simplified model is maintained with an emphasis on the biospecimen provenance. This structure provides a series of entities for which key metadata can be attached across both single cell and spatial assays, despite the differences in methodologies.

Following the creation of the assay-agnostic biospecimen provenance diagram, a consensus was reached on a core set of data elements across the library construction lifecycle. Experts from fourteen organizations, including biomedical researchers, experimenters, data analysts, and informaticians, contributed to the definition of data elements for each stage of the library construction process (see **Supplemental Table S3** for comprehensive list of donor, specimen, and library metadata). The model entities and their associated metadata properties were implemented in the NIMP portal, and continue to be updated with operational fields, quality metrics and steps, and additional fields to support submission tracking and data release. The fields represented here focus on those required for scientific and computational use of sequencing data; administrative and operational fields used for logistics such as shipping information, sample tracking and laboratory operations are not described in this model. The portal supports sequencing library submission and associated metadata management for entities and processes from tissue to library pool.

## 4: Harmonized Biospecimen Metadata Schemas

Human and non-human primate tissue acquisition and sharing is a critical component of the BICAN consortium. While the NIH NeuroBioBank (RRID:SCR_003131) is the major source for human tissue in BICAN, individual laboratories may also source human tissues from heterogeneous sites, and non-human primate and mouse tissue from a variety of laboratories and sources. This heterogeneity necessitated a metadata framework that supports capture of provenance across multiple sources, species, and downstream uses. The NIMP metadata portal supports provenance tracking across tissue sources, species, specimen types and downstream uses.

Developing donor and tissue metadata standards in this environment presented consortium-specific challenges that shaped our approach. Different species require distinct metadata fields; for example, human donors have associated clinical and demographic information, whereas colony-housed non-human primates may social and behavioral observations. Metadata availability also varies by tissue source, such as biobank versus laboratory source, and by developmental stage. In addition, tissue preparation requirements differ by assays; Patch-seq experiments require live tissue experiments whereas molecular profiling often utilizes frozen or postmortem tissue.

Rather than enforcing a rigid, maximal schema, we adopted an inclusive harmonization strategy that accommodated unique features of diverse tissue sources. Our approach documented both requested and available metadata fields and included iterative mapping and harmonization of data elements across a distributed set of laboratories and tissue banks. For human donors, initial requirements were developed in collaboration with the BICAN Specimens Working Group and aligned with NeuroBioBank (“The NIH NeuroBioBank” 2018), the primary brain tissue source for BICAN, particularly given its mature metadata schema. These requirements were then reviewed by stakeholders to accommodate donor tissue obtained from other sources. This alignment allowed BICAN-dedicated human brain collections to be maximized for incorporation of available donor metadata from NIH NeuroBioBank and other dedicated BICAN brain banks (Berry et al. 2025). These metadata included a comprehensive list including general subject fields, medical and clinically relevant fields, among other donor-specific information (**Fig. 3B**), as well as fields regarding the availability of antemortem or postmortem MRI imaging general specimen fields with information about RIN values and other data that reflect tissue quality or cellular composition.

**Figure 3.**
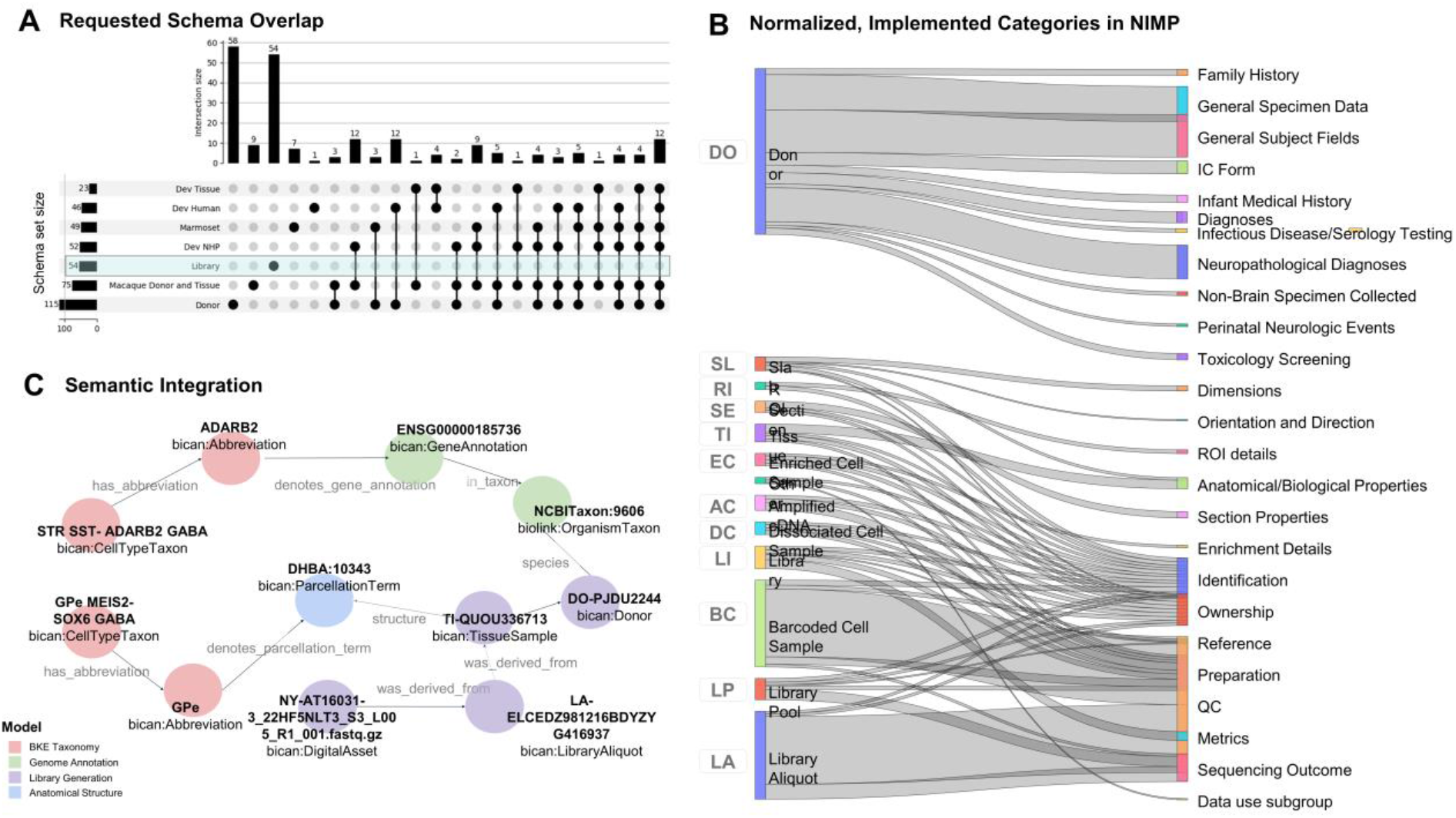
Donor, Tissue, and Library Metadata Schemas. Consortium metadata schemas were integrated to map and uniquely identify metadata elements entering the ecosystem for diverse species, developmental stages and techniques, covering basic donor/subject data, tissue or specimen data, and sequencing-related data. **A, the** upset plot shows the number of elements per schema on the bar plot on the left; the overlap of elements of these schemas is shown as subsets in the column chart across the top with black dots below which schemas share those fields. Note that the library metadata schema represents a non-over-lapping set of elements. **B**, Sankey diagram with the final implemented schemas at NIMP that includes properties mapped to resource categories generally associated with the biospecimen entities from Figure 2. Specimen-type icons correspond to the biological entities Figure 2, providing a direct visual mapping between abstract workflow stages and implemented metadata categories. **C**, Example semantic graph enabled by LinkML models and the *bkbit* Python library. Domain-specific data schemas (e.g., cell types, tissue samples, genome annotations, and anatomical structures) are designed to interoperate as a connected knowledge graph (KG) via schema-compliant JSON-LD representations.

The prioritized use cases were the following. First, the schemas should enable a specimen request workflow for centralized banked human tissue by describing donors and sequential steps of tissue processing, while also supporting multiple species in a common framework (**Fig. 1B**). Second, they needed to support biospecimen provenance records for the core sequencing workflow, including metadata about intermediate biospecimens generated during sequencing library construction.

The initial outcome of this process was the development of multiple schemas, individualized for the diverse donor and tissue types for which laboratories requested metadata support, including human donor, developmental tissue, and rhesus macaque and marmoset; see **Fig. 3A**). Because these metadata needs emerged at different times and from different groups, these schemas were developed through a staggered harmonization process rather than as a single centralized effort. For each schema, data dictionaries were created to define each metadata element, including descriptions, data types, controlled vocabulary requirements, and related mappings. Metadata elements within each schema were mapped to a single index with BICAN UUIDs, so that fields that exist in multiple schemas are identifiable as the same item. In contrast, sequencing library metadata elements were developed through a more centralized process and were consolidated into a single list of fields through a dedicated task force effort. Together, these schemas, data dictionaries, UUID-indexed mappings, and library metadata fields provide a record of how each group’s metadata schemas had been harmonized into the ecosystem.

The NIMP portal supports collection of the appropriate metadata per donor and tissue type with flexibility per tissue type or source while enforcing requirements for core fields (**Table S3**). BICAN-developed models were designed to be interconnected and interoperable. Entities across different domain-specific models (e.g., donors, anatomical structures, and library pools) are linked through formally defined semantic relationships. The schema-compliant JSON-LD we generate explicitly encodes cross-schema links and can be represented as an RDF graph (**Fig. 3C**). This design ensures that donor and tissue metadata are not isolated descriptors but are first-class, queryable components that can be represented in a knowledge graph.

A central tenet of BICAN is rapid and open data sharing, and the strategies for collecting and distributing BICAN data and associated metadata are intended to provide data access as openly as possible while respecting consent, privacy, and data use constraints. This approach required careful consideration of issues around patient consent and de-identification. Two key metadata policies were adopted to de-risk data releases. First, only de-identified metadata should be entered into the BICAN data ecosystem. Second, fields that could present privacy risks despite presumed de-identification were identified for additional review and marked to be withheld from public release (**Table S3**).

## 5: Cell Annotation and Taxonomy Standards

A major goal of BICAN is to produce a reference standard for brain cell types and their relationships that enables the neuroscience community to integrate multimodal data and knowledge into a common framework. Standardized schemas for representing cell type taxonomies are therefore a critical prerequisite for this integration effort.

Cell type taxonomies provide a structured way to represent cell types or provisional cell type classes, their hierarchical relationships, and the evidence used to define and refine them. In recent brain cell atlas studies, cell types are first organized into broad classes based on anatomic location and major neurotransmitter families. Disputes in cell type identification and nomenclature may arise when different methodologies to phenotype neurons cannot be easily reconciled, such as using morphology, electrophysiology, or connectivity in individual assays. This results in a patchwork of identified cell types with inconsistent naming and incomplete feature sets.

Cell type taxonomies in BICAN were built using a data-driven approach. On the premise that cell types with distinct functions should be discriminable on the basis of their gene expression, single-cell transcriptomic, epigenomic and multiomic data were analyzed and hierarchically clustered to create a taxonomy of cell types. Many transcriptomically defined cell populations show high conservation across individuals and species indicating that they represent real biological diversity. However, separating functionally distinct cell types from variation due to cell state or biological gradients, and refining boundaries among closely related cell types requires the integration of additional data types and prior knowledge. Multimodal data and expert annotation are therefore layered onto molecular taxonomies to provide context, support interpretation and cell type naming, evaluate cluster coherence, and to organize cell types into hierarchical groupings such as classes, subclasses, and types. Taxonomies created from a single assay type, dataset or species may also require alignment and integration with data derived from other modalities and species. The final output is a set of putative hierarchical cell types supported by quantitative and qualitative evidence linked to the underlying molecular data. This evidence, together with the source data, annotations and taxonomy versions, is critical to capture and track as provenance.

Hierarchical, data-driven cell type taxonomies were exemplified by early single-cell transcriptomic studies such as Tasic et al. 2016 (Tasic et al., 2016) **(Fig. 4F**), and have been widely adopted in collaborative cell-typing efforts. As these taxonomies have expanded to include additional annotations, evidence and mappings, a standard structure to represent this information was needed to support the basic premise of data-driven cell type taxonomies while remaining extensible to support new data types and annotations. This need is especially important because cell type taxonomies are no longer merely static reference products; they are increasingly used as computable resources that support taxonomy building, sharing, mapping, and community annotation. The AIT format described here enables the creation of tools for building a taxonomy based upon any set of high quality single-cell omics data with computational and manual processes for analysis and annotation; thus, other scientists can create or use cell type taxonomies in AIT format with relative ease. Four specific use cases for the creation and usage of a cell type taxonomy schema are shown in **Fig. 4A-D**.

**Figure 4.**
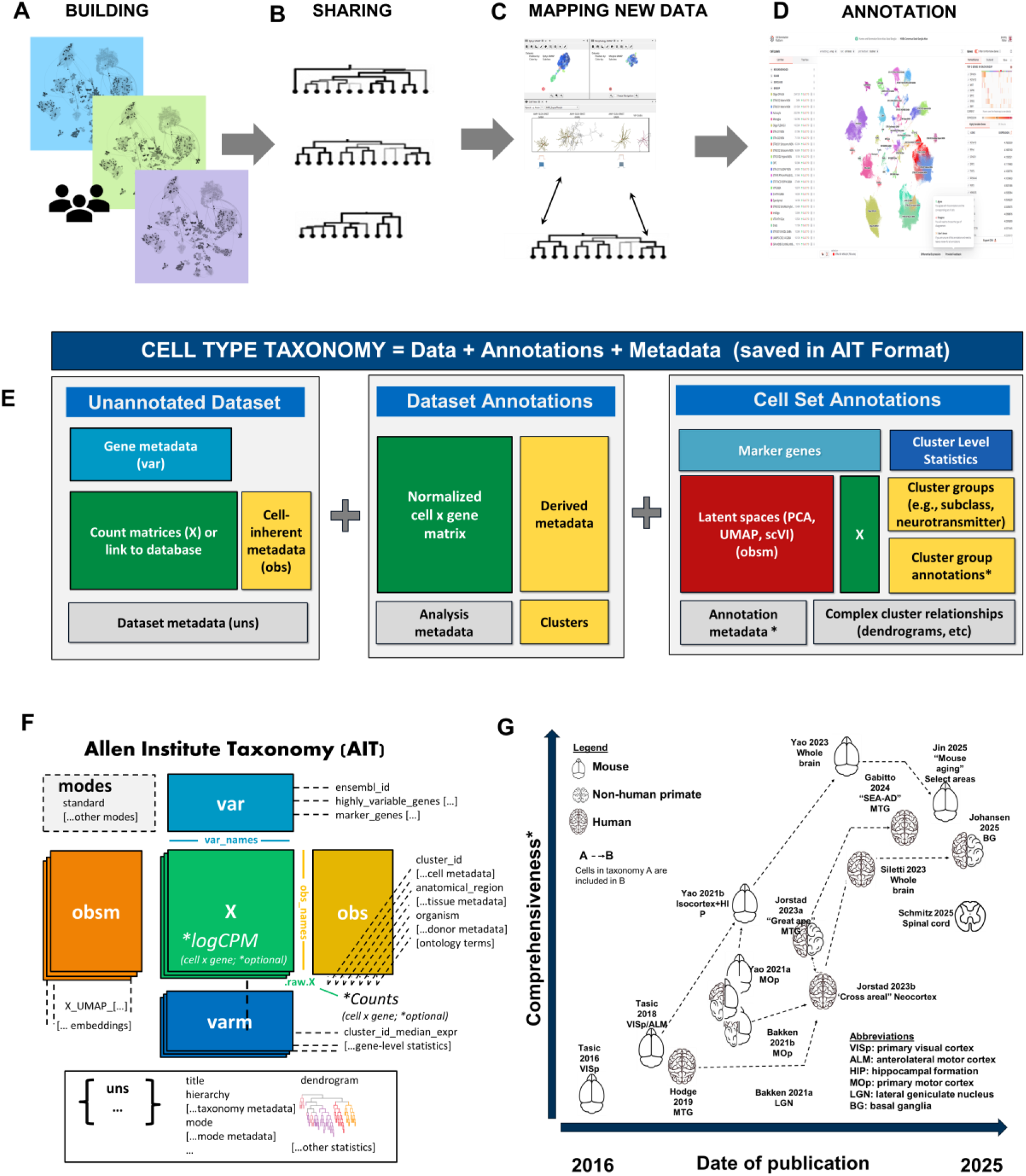
Allen Institute Taxonomy (AIT) Schema. The AIT schema standard was developed to support four main uses cases (**A-D**). E, The AIT format schema format extends the widely adopted CELLxGENE schema by modularizing the constituent parts and adds “modes”. The taxonomy becomes a combination of unannotated data, dataset annotations, and cell set annotations (**E**). The taxonomy then manages these as different modes **(F**). Taxonomies generated from 2016-2025 have been re-released in this format to enable access in a consistent way (**G**).

First, researchers can assemble data, metadata, clustering results and annotations into a taxonomy (**Fig. 4A**). The scrattch.taxonomy library (RRID:SCR_027910; https://alleninstitute.github.io/scrattch.mapping) provides a single function that creates an AIT file for users when provided basic single-cell transcriptomics data and metadata files (buildTaxonomy), along with other functions for standardizing metadata, predicting ontology terms from text values, and defining taxonomy modes.

Second, encoding taxonomies into a common single-file format enables broad sharing of these taxonomies (**Fig. 4B, F**). More than a dozen taxonomies generated between 2016 and 2025 have been re-released in this common format (T. E. Bakken, Jorstad, et al. 2021; T. E. Bakken, Van Velthoven, et al. 2021; Gabitto et al. 2024; Hodge et al. 2019; Jin et al. 2025; Johansen et al. 2025; Jorstad, Close, et al. 2023; Jorstad, Song, et al. 2023; Schmitz et al. 2026; Siletti et al. 2023; Tasic et al. 2016, 2018; Yao et al. 2023; Yao, Van Velthoven, et al. 2021; Yao, Liu, et al. 2021). A user can now explore any of a decade’s worth of related brain cell type taxonomies encoded in a common format, and use with established tools for omics data visualization and exploration (e.g., CZI CELLxGENE).

Third, a common format enables users to map their own data to these taxonomies (**Fig. 4C)**. MapMyCells (RRID:SCR_024672) provides a highly robust and user-friendly interface for assigning cell types from select BICAN taxonomies to user-provided single-cell RNA-seq data (Daniel et al. 2026). The scrattch.mapping library extends this functionality to any cell type taxonomy saved in AIT format, providing flexibility in aligning user data to any public cell type taxonomy.

Finally, the AIT taxonomy file format is closely aligned to the schema for Cell Annotation Platform (Cell Annotation Platform (CAP); RRID:SCR_022797), allowing transformation and cross-platform ingest to support community annotation, including suggestions for updates to cell type nomenclature, adjustments to cluster resolution, and inclusion of supporting literature (**Fig. 4D)**. These annotations can then also be added as layers back into the flexible AIT format to support visualization of additional cell type features.

Together, these use cases define the requirements for the AIT schema: the format must preserve molecular data and metadata, represent hierarchical cell type annotations, support multiple views of a taxonomy and capture the evidence and provenance needed for comparison, reuse, and ongoing refinement. The Allen Institute Taxonomy (AIT) schema defines a structured and compartmentalized standard for storing and distributing all components of a curated cell type taxonomy within an extensible AnnData (Virshup et al. 2024) framework (**Fig. 4 E**), including definitions for terms such as “taxonomy”, “dataset”, “annotation”, “metadata”, and “data”. Building upon the widely adopted CELLxGENE schema, AIT extends its components to accommodate the specific needs of the BICAN consortium and the broader community developing large-scale cell type taxonomies (as discussed below). The format encapsulates every conceptual layer of a taxonomy including single-cell data, metadata, hierarchical annotations, and computational tooling within a formalized schema that provides consistent, well-defined fields for accessing each component, addressing the inconsistencies that have characterized prior taxonomy releases. The schema adopts normative keywords (“MUST,” “RECOMMENDED,” etc.) to enforce reproducibility and alignment across implementations. Finally, to support the formalized schema and controlled vocabulary, the AIT framework currently includes R-based workflows to generate and work with h5ad files following the AIT schema (**Fig. 4 E**).

AIT extends the CELLxGENE and related schema in a few key ways. First, the schema itself includes terms specific to the cell type taxonomy usage that are not included in broader data format standards. These include fields specific for cell type hierarchies, as well as precomputed values and statistics that facilitate cell type mapping and other analyses. Second, the AIT format is designed to flexibly accommodate taxonomy ‘modes’ representing subsets of cell types, such as all neuronal types, all cells from a specific brain region, or genes, such as a spatial transcriptomics gene panel. Taxonomy modes provide flexibility to perform many different analyses using the same base taxonomy, rather than needing to save multiple copies corresponding to different subsets of cells from the same experiment. Finally, we have integrated the Cell Annotation Schema (CAS; RRID:SCR_027909) into the AIT schema. CAS has gained traction through its compatibility with widely adopted platforms including CZI CELLxGENE and Cell Annotation Platform and can be used as a more robust method of storing cell type annotations than the more commonly used Google Sheets. By embedding CAS within the unstructured (*uns*) component of AIT files (**Fig. 4F**), we enable extended annotation capabilities—including ontology mappings, evidence tracking, and marker expression documentation—without disrupting existing data structures, through strategic nesting of information. By standardizing the structure and terminology of taxonomies, AIT promotes transparency, version control, and interoperability, enabling reliable comparison and reuse of taxonomies in the emerging era of comprehensive, whole-brain cell type atlases.

The AIT schema also supports interaction with formal ontology development by organizing provisional, data-driven taxonomies in ways that can inform community cell type representation. The Cell Ontology (CL) (Tan et al. 2026; Diehl et al. 2016) was developed to provide a structured representation of cell types that can allow cross-modality comparison where the data supports it. However, while CL holds curated and approved cell types, identification of new cell types at unprecedented granularity from single-cell omics technologies has challenged real-time CL content creation. To overcome this bottleneck, an approach for defining cell types in CL that combines the species and anatomic structure source, and parent cell type already represented in the CL, and the optimal combination of marker genes was proposed(T. Bakken et al. 2017). Thus, robust mechanisms for storing key metadata, data, and knowledge, including taxonomic relationships, anatomic structure, and marker gene information has become critical for driving CL development (Tan et al. 2026, 2023). Indeed, the hierarchical structures captured in the AIT align with the directed acyclic graphical structures in CL, facilitating parent-child cell type designation. Thus, the structures and metadata captured in the AIT are valuable for the representation of provisional and new cell types in the CL (Tan et al. 2023).

Together these examples show the value of packaging cell type taxonomies in a single file, because tools such as CELLxGENE and scrattch.mapping rely on complete.h5ad files that package data, metadata, and annotations together in a common structure. However, as atlas scale datasets continue to grow, it may becoming increasingly important to separate large data from taxonomy components and to compartmentalize all components to avoid the need to download, open, or upload huge and unwieldy files. As one example, h5ad files that include data matrices with more than a few hundred thousand cells cannot be opened and therefore are unusable in R. AIT partially addresses this by making the data frames optional, but this concept of modularity will become increasingly important as data sets increase in size.

The AIT schema accommodates both paradigms: it supports comprehensive single-file packaging for immediate usability (AIT) while providing clear separation of concepts (**Fig 4G**) that facilitate use by tools like Allen Brain Cell Atlas and MapMyCells. Rather than prescribing a rigid final structure, our framework establishes organizing principles and field mappings that can adapt as individual standards mature and, ideally, converge toward a unified community standard.

## 6. Discussion

Standards constitute the essential infrastructure that enables systematic scientific inquiry, reproducible research, and cumulative knowledge advancement. In neuroscience, the diversity of experimental approaches, data types, and analytical methods that have been employed to understand cell type diversity present a data integration challenge. To address these challenges, we engaged in careful assessment, design, curation, and standardization of data, metadata, and knowledge for cells, donors, tissues, and associated laboratory information using a consensus-based development process. The standards described here establish a multi-layered framework for connecting biospecimen provenance, donor and tissue metadata, sequencing metadata and cell type taxonomy knowledge within the BICAN ecosystem.

Our three examples, the biospecimen provenance model for single-cell and spatial omics data, cross-species donor harmonization, and cell type taxonomy, reveal common challenges that transcend specific technical domains: reconciling redundant information across groups, documenting transparent decision-making, fitting standardization alongside ongoing scientific operations, and balancing interoperability with metadata fidelity. These tensions are neither unique to BICAN nor isolated incidents but represent fundamental challenges inherent in standardizing complex, evolving data across heterogeneous research environments. The solutions we developed (sustained Working Groups and targeted Task Forces, structured curation pipelines from collaborative spreadsheets to machine-actionable LinkML models, and version-controlled public repositories) provide reusable templates applicable across biomedical consortia.

A key contribution of this work is the use of the biospecimen provenance as a practical abstraction for harmonizing diverse experimental workflows. We were able to standardize core metadata associated with all biological samples tracked in this large, multi-institutional project by establishing consensus across the consortium on the use of the NIMP portal for tissue registration and the application of controlled vocabulary terms. This agreement ensured that key attributes of donor tissues and derived biosamples were captured in a consistent, structured manner at the point of entry, reducing ambiguity and variability introduced by laboratory-specific conventions. As a result, tissue provenance, processing history, and biological context could be reliably linked to downstream single-cell and spatial omics data, enabling seamless integration with the unified sequencing metadata framework. The integration of partner data archives provides additional reinforcement of standards usage through requirement of appropriate identifiers and ingest of data directly from the NIMP portal. Collectively, this approach demonstrates how centralized sample registration coupled with consensus-based semantic standards substantially improves metadata quality, traceability, and analytical rigor in large consortium-scale biomedical studies.

The technical infrastructure we established transforms human-accessible documentation into computational resources that enable validation, transformation, and semantic interoperability. Our pipeline from metadata spreadsheets through LinkML schemas to provenance-aware JSON-LD, complemented by the process diagrams, bridges the gap between collaborative consensus-building and the computational requirements of modern data science. This infrastructure represents a useful template to support the initiation of a new metadata schema standardization project. Importantly, the schemas provide a framework for recording provenance, identifiers, and relationships, but complete provenance capture depends on consistent implementation, validation, and curation within operational workflows.

BICAN metadata standards support FAIR principles as foundational infrastructure that extends beyond our consortium. Our commitment to FAIR principles includes ensuring data are “Findable” through persistent identifiers and structured metadata, “Accessible” by publishing data at archives, “Interoperable” through community-standard ontologies, and “Reusable” through comprehensive documentation and open licensing (Wilkinson et al. 2016). These principles apply to both primary data and to derived knowledge products such as cell type taxonomies, annotations, and evidence records.

Cell type taxonomies represent an important class of evolving knowledge products. In the case of the cell type taxonomy and annotation standard, the need to share data and knowledge together requires that tools can create a data package, map to the data package, and visualize the data package. Notably as part of the recent BICAN efforts, consensus cross-species cell type taxonomies were released for the basal ganglia (Johansen et al. 2025), alongside the MapMyCells mapping tool (Daniel et al. 2026). The AIT schema supports these needs by packaging molecular data, metadata, hierarchical groupings, supporting evidence and provenance in a computable format that can be used for taxonomy building, sharing, mapping, visualization, and community annotation.

However, the rapid pace of evolution of our knowledge about cell types introduces additional challenges. Cell type taxonomies derived using this data-driven approach will necessarily change based on the addition of new data, both in scale and in modality, as well as new analysis and annotation layers. The AIT format can capture the data, evidence, and annotations in layers, and the use of Cell Ontology to document cell types with their molecular evidence provides a supporting formal record after mature analysis. However, managing the relationships between cell types observed in different experiments, taxonomies, or taxonomy versions will require additional attention. Future models will need to capture how cell types overlap, diverge, merge, split or otherwise change between taxonomies; how relationships across species are inferred and qualified; and how data-driven cell type definitions evolve in conjunction with classical nomenclature.

AIT highlights a wider infrastructure tension between self-contained file formats and modular knowledge systems. While the scientific community continues to rely heavily on self-contained file formats for sharing, analysis, and visualization, distributed storage and database-driven architectures represent the future of cell atlas efforts, and both tightly integrated and highly modular approaches are needed. Self-contained.h5ad files support easier sharing and reuse, because all necessary information is indexed and packaged for ready transformation for use in a variety of tools. The popularity of this format echoes advances made for sharing neuroimaging data using the BIDS format (Gorgolewski et al. 2016) and for sharing full neurophysiology experiments with NWB (Rübel et al. 2022). However, atlas-scale data will increasingly require separation of molecular matrices, potentially composed of hundreds of millions of cells measured across tens of thousands of genes or features, from the metadata, annotations, evidence and derived summaries. AIT supports modularity and currently makes the data matrices optional to allow for a lightweight version of the taxonomy to be shared, while preserving conceptual organization.

Critical to successful standards adoption is the establishment of governance frameworks that address long-term maintenance, version control, and conflict resolution. Collaborative, time-bounded efforts often experience a tension due to synchronous development of metadata standards with implementation. Experimental workflows need to be unblocked requiring early deployment of draft metadata schemas prior to full integration and harmonization, and continue to require evolution to respond to emergent issues. This creates a risk of discrepancies between the official standard and current implementation. However, early implementation also provides real-world validation prior to full standards development. Iteration becomes a necessity and a feature. Governance frameworks help support the initial creation and evolution of standards, and require collaboration between the subject matter expert experimentalists, data scientists, and engineers supporting the infrastructure. A potential pitfall for standards can be the distance that lies between an idealized standard and real-world implementations. A standard must be usable to be useful, and that often necessitates that there are not only use cases for it, but toolchains that incorporate those standards.

Another key motivator for the development and cross-consortium adoption of well-documented data models and metadata standards is the proliferation of AI models and tools that are being developed to support data analysis and experimental design within the broader research community. High-quality, well-documented, and expertly-curated datasets and knowledge representations such as those generated by the BICAN consortium are critical for building and validating such models, and the standards framework we describe in this study is a key component in the BICAN ecosystem for facilitating the use of consortium datasets by AI developers.

The success of BICAN’s standards development process demonstrates that addressing questions exceeding individual laboratory capacity requires coordinated effort transcending traditional organizational boundaries. By embracing rather than resisting consortium complexity, developing governance structures balancing efficiency with inclusivity, and investing in metadata infrastructure as prerequisite rather than afterthought, we create lasting resources that transform isolated investigations into coherent, comparative scientific understanding. The insights we provide—documenting both successes and difficulties in achieving consensus, describing technical pipelines connecting human collaboration to computational infrastructure, and establishing governance frameworks for sustained community engagement—offer practical guidance transferable across biomedical consortia navigating similar standardization challenges. This long-term value of these standards lies not only in the specific schemas described here, but in the framework they establish for maintaining consistent identifiers, provenance, evidence and relationships across data and knowledge products in a community-wide manner.

## Limitations

While we have captured many essential components required to ensure FAIRness of consortium datasets, there are gaps in our current documentation and practices that can be addressed. Experimental protocols for bioinformatics workflows include a significant computational component, and with the importance of statistical and machine learning methods in these workflows, simply providing links to code repositories and plaintext protocols may not be adequate to ensure reproducibility. The ability to accurately reproduce complex analyses requires access not only to static code, but also reproduction of the programming environment, including versions and dependencies of any packages used and documentation of critical environment variables. Furthermore, analytical workflows that include a significant probabilistic component such as cell type clustering can require documentation of the exact algorithmic parameters used to generate outputs for a given instance of an analysis run. While containerization of computational components helps address the reproducibility objective, it does not provide for the standardized description of the process.

Additional limitations arise from the scope and implementation context of the standards described here. The schemas reflect BICAN operational priorities and may require adaptation for other consortia or scientific topics. The publicly available scientific metadata models here also do not capture all administrative, operational, and logistical fields used internally. The biospecimen provenance model represents a high level abstraction to enable metadata capture, and may not represent literal experimental steps. Finally, cell type taxonomy standards remains an active area of development, and additional mechanisms are needed to capture the dynamic relationships between species, modalities, versions, and releases.

## Resource Availability

Requests for further information and resources should be directed to and will be fulfilled by the lead contact, Carol Thompson.

All original code and standards documentation has been deposited in GitHub as detailed below and is publicly available as of the date of publication. BICAN metadata schema and AIT standards documentation are available on GitHub (https://github.com/brain-bican/metadata-schemas), and specific release versions have been created with Zenodo (https://doi.org/10.5281/zenodo.19377945). BICAN LinkML models are available on GitHub (https://github.com/brain-bican/models). Accessibility is ensured through open licensing (CC BY 4.0) permitting unrestricted use, modification, and redistribution with attribution.

The scrattch package is available on GitHub (https://alleninstitute.github.io/scrattch.mapping).

Implemented metadata fields can be viewed with the NIMP terminology browser (https://brain-specimenportal.org/terminology_browser).

Existing cell type taxonomies provided in the AIT format can be found at https://brain-map.org/our-research/cell-types-taxonomies.

## Acknowledgements

This publication was supported by and coordinated through the BRAIN Initiative Cell Atlas Network (BICAN). This work was funded by the Allen Institute and by the National Institutes of Health awards U24MH130919 (MH), U24MH130918 (SM), U24MH130968 (OW), R24MH114788 (OW), R24MH114793 (AR), U24MH130988 (GQZ), UM1MH130981 (EL), UM1MH130966 (CV), UM1MH130991 (HH), and U24NS133077 (JM). This research was also supported in part by the Intramural Research Program of the National Institutes of Health (NIH). The contributions of the NIH authors are considered Works of the United States Government. The findings and conclusions presented in this paper are those of the authors and do not necessarily reflect the views of the NIH or the U.S. Department of Health and Human Services. This research was funded in part by Wellcome (grant no. 220540/Z/20/A, “Wellcome Sanger Institute Quinquennial Review 2021–2026”). We thank the founder of the Allen Institute, Paul G. Allen, for his vision, encouragement and support.

## Declaration of Interests

Dr. Martone is a co-founder and serves on the board of SciCrunch, Inc. The terms of this arrangement have been reviewed and approved by the University of California, San Diego in accordance with its conflict of interest policies. Dr. Bandrowski is a co-founder and current CEO of SciCrunch, Inc. The terms of this arrangement have been reviewed and approved by the University of California, San Diego in accordance with its conflict of interest policies.

Declaration of generative AI and AI-assisted technologies in the writing process: During the preparation of this work, the author used ChatGPT in order to assess manuscript clarity and flow. After using this tool, the author reviewed and edited the content as needed and takes full responsibility for the content of the published article.

## STAR Methods

### Key resources table

| REAGENT or RESOURCE | SOURCE | IDENTIFIER |
| --- | --- | --- |
| Software and algorithms |  |  |
| NHASH Identifiers | BICAN: Neuronanatomy Informed Metadata Portal (NIMP). UTHealth, Houston | RRID:SCR_025313 |
| scrattch package | Allen Institute | https://alleninstitute.github.io/scrattch.mapping |
| Other |  |  |
| Research Resource Identifiers | SciCrunch | RRID:SCR_015949 |
| Harmonized Ontology of Mammalian Brain Anatomy (HOMBA) | BICAN; Allen Institute, Brain Science | RRID:SCR_027628 |
| Cell Ontology |  | RRID:SCR_004251 |
| Atlas Ontology Model |  | RRID:SCR_023813 |
| Ontology of Biomedical Investigations |  | RRID:SCR_006266 |
| BICAN Metadata Standards | BICAN; Allen Institute, Brain Science | doi: 10.5281/zenodo.19377944 |
| Cell Annotation and Taxonomy Standard (CAS) | BICAN; Allen Institute, Brain Science | doi: 10.5281/zenodo.19377944 |
| BICAN Knowledge Base LinkML Schemas | BICAN; MIT | https://github.com/brain-bican/models |
| Allen Institute Taxonomy Standard (AIT) | BICAN; Allen Institute, Brain Science | RRID:SCR_027629 |
| NIMP terminology browser | BICAN; UTHealth | https://brain-specimenportal.org/terminology_browser |

