## Supplemental Figure S1 for "A layered standards framework for integrating single-cell and spatial omics data into brain cell atlases"

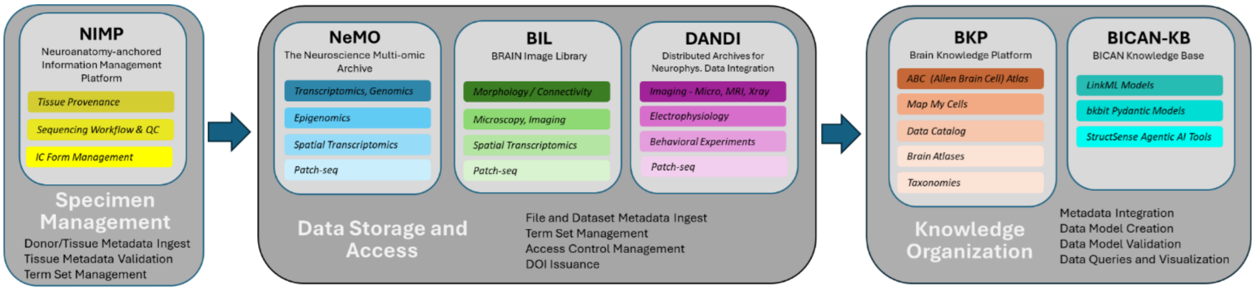


**Figure S1. Overview of the BICAN Data Ecosystem and standards-enabled data flow.** The BICAN ecosystem integrates specimen management, multimodal data storage, and knowledge organization through coordinated infrastructure and metadata standards. Biological specimens and associated metadata are registered in the Neuroanatomy-anchored Information Management Platform (NIMP) which manages consortium specimen requests, tracking of tissue provenance, sequencing status and quality control, and management of institutional certification (IC) forms that document permissions for the datasets. The data are deposited into domain-specific archives, including Neuroscience Multi-Omic (NeMO) Data Archive, Brain Image Library (BIL), and Distributed Archives for Neurophysiology Data Integration (DANDI), where dataset-level metadata, identifiers and access controls are managed. These data are subsequently integrated into downstream knowledge platforms, including the Brain Knowledge Platform (BKP) and BICAN Knowledge Base (BICAN-KB), which support data model creation, validation, integration, and user-facing tools. This ecosystem enables structured flow of data from specimen to knowledge, supporting interoperability, reuse and analysis across the consortium and research community.
