## Supplemental Table S1 for "A layered standards framework for integrating single-cell and spatial omics data into brain cell atlases"

**Table S1. Selected BICAN-developed standards, schemas, identifiers, community ontology contributions, and interoperability targets.**

| Standard (RRID/PID) | BICAN Usage | Description |
|---|---|---|
| ***Identifiers*** | | |
| **NHash Identifier**<br>RRID:SCR_025313 | • Developed by BICAN<br>• Issued for NIMP-managed entities | Light-weight, block chain style resource identifiers for tracking research resource linkage, provenance, utilization, and visualization. NHash identifiers incorporate a set of cryptographic hash functions based on N-grams, with combination of additional encryption techniques such as shift cipher. |
| **Research Resource Identifiers**<br>RRID:SCR_015949 | • Used by BICAN<br>• Issued for major resources | Identifier repository for research resources. |
| ***Ontologies*** | | |
| **Harmonized Ontology of Mammalian Brain Anatomy (HOMBA)**<br>RRID:SCR_027628 | • Developed by BICAN<br>• Used to annotate BICAN tissue, data and cells | Harmonized cross-species taxonomy of 2341 brain and spinal cord structures. Derived from the Allen Developing Human Brain Atlas (DHBA) ontology, the HOMBA is hierarchical, allowing users to aggregate structures from fine grain parcellations to broad regions. Terminology is harmonized across human, primate, and rodent structures with synonymous terms and includes transient developmental structures. |
| **Cell Ontology**<br>RRID:SCR_004251 | • Updated by BICAN to include new cell types<br>• Includes new BICAN-derived basal ganglia cell types | A reference ontology designed to classify and describe cell types across different organisms. It serves as a resource for model organism and bioinformatics databases. |
| **Atlas Ontology Model**<br>RRID:SCR_023813 | • Used by BICAN<br>• Packaging of brain common coordinate frameworks | A standardized, machine-readable framework designed to harmonize brain atlases by defining the relationships between four core elements: reference data, coordinate system, annotation set, and terminology |
| **Ontology of Biomedical Investigations**<br>RRID:SCR_006266 | • Used by BICAN<br>• Integrated with LinkML models | An integrated ontology for the description of life-science and clinical investigations |
| ***Metadata*** | | |
| **BICAN Metadata Standards**<br>doi: 10.5281/zenodo.19377944 | • Developed by BICAN<br>• Documents metadata schemas | Full suite of BICAN metadata standards hosted in GitHub including human donor, developmental tissue, and non-human primate schemas. This repository also hosts the Allen Institute Taxonomy and Cell Annotation Standards. |
| **Cell Annotation and Taxonomy Standard (CAS)**<br>doi: 10.5281/zenodo.19377944 | • Developed by BICAN<br>• Used for cell type annotation | The cell annotation and taxonomy standard establishes a unified framework for identifying and classifying cell types across datasets. This standard incorporates hierarchical cell type ontologies, marker gene specifications, and confidence metrics for cell type assignments. The schema supports both automated and manual annotation workflows, enabling consistent cell type nomenclature across research groups and experimental platforms. |
| **BICAN Knowledge Base LinkML Schemas**<br>https://github.com/brain-bican/models | • Developed by BICAN<br>• Used for BICAN data models | Brain data models in LinkML for the development of the Brain Cell Knowledge Base (BCKB) to ingest and standardize comprehensive cell type information from BICAN's development of a multimodal, multi-species brain cell atlas and disseminate that atlas as an open and interactive community resource for advancing knowledge of the brain |
| ***Data Formats*** | | |
| **Allen Institute Taxonomy Standard (AIT)**<br>RRID:SCR_027629 | • Documented by BICAN<br>• Used for cell type taxonomies | Aligned taxonomy format and supporting tools. |
| **Brain Imaging Data Standard (BIDS)**<br>RRID:SCR_016124 | • Used by BICAN<br>• MRI data | Provides a consistent way to organize your neuroimaging data |
| **SWC**<br>RRID:SCR_027631 | • Used by BICAN<br>• Morphology data at BIL data archive | Text-based (ASCII text) files that describe three-dimensional neuronal or glial morphology. These digital reconstructions represent morphology as a vectorized tree structure, made of a series of connected nodes. |
| **Neurodata Without Borders (NWB 2.0)**<br>RRID:SCR_015242 | • Used by BICAN<br>• Patch-seq electrophysiology data at DANDI | A data standard for neurophysiology to share, archive, use, and build analysis tools for neurophysiology data. |
