## Supplemental Table S2 for "A layered standards framework for integrating single-cell and spatial omics data into brain cell atlases"

| Step / Method | Tissue Sectioning | Sample prep / isolation | Optional Enrichment | Barcoding / tagging | Other Chemistry Steps | Imaging | Amplification | Adapter ligation & library completion | Methods References |
|---|---|---|---|---|---|---|---|---|---|
| **Single cell transcriptomic and multiomic assays** | | | | | | | | | |
| | | TI → DC | DC → EC | EC → BC | na | na | BC → AC | AC → LI | |
| **Multiome (10x ATAC + GEX)** | na | Nuclei extraction from cells/tissues. | Optional FACS sorting to enrich cell populations | GEM encapsulation with barcoded beads capturing both mRNA and ATAC fragments (Droplet barcodes). | mRNA Capture — Reverse transcription in droplets with oligo-dT; ATAC — Tn5 transposition of open chromatin. | na | Pre-amplify cDNA & ATAC fragments; THEN split for library prep. | Illumina adapters + sample indices added during library PCR (for ATAC and GEX). | https://www.10xgenomics.com/blog/introducing-chromium-single-cell-multiome-atac-gene-expression |
| **snm3C-seq** | na | Nuclei extraction from cells/tissues. | Single nuclei sorted into plates with optional enrichment. | Cell barcode introduced (plate indexing). | Chromatin conformation capture + bisulfite conversion for methylome + proximity ligation for contacts. | na | PCR amplify bisulfite-converted contact/methylome fragments. | Adapters are part of PCR products ready for NGS. | Luo et al. (2019). *Nature Methods 1* 6 (10) 999-1006. |
| **snmCT-seq** | na | Cell / Nuclei extraction from cells/tissues. | Single nuclei sorted into plates with optional enrichment. | Cell barcode introduced (plate indexing). | RNA reverse transcription + bisulfite conversion for methylome without physical DNA/RNA separation. | na | PCR amplify barcoded mixed RNA/methylome molecules to build sequencing libraries. | Adapters included in amplified products, ready for sequencing. | Luo et al. (2018) *bioRxiv* 434845; doi: https://doi.org/10.1101/434845 |
| **Droplet Paired-Tag** | | Nuclei extraction from cells/tissues. | Optional FACS sorting to enrich nuclei | Combinatorial indexing of histone marks and transcripts with cell barcode (Droplet barcodes ). | Capture of histone modification targets (antibody-Tn5 chemistry) + mRNA capture inside droplets. | na | Amplify both transcript and epigenetic (histone) fragments post-barcoding. | Sequencing adapters added during PCR/ligation steps. | Xie et al. (2023) *Nature Structural & Molecular Biology* 30, 1428-1433. |
| **Sequencing-based spatial transcriptomic assays** | | | | | | | | | |
| | TI → SE | SE → DC | DC | EC → BC | | | BC → AC | AC → LI | |
| **DBiT-Seq** | Tissue sectioned | Tissue mounted on slides | Incubation with antibody-derived DNA Tags for proteins of interest. | Two sets of orthogonal spatial barcodes delivered before or after RT via microfluidic channels, ligated in situ. | In situ reverse transcription of mRNA to cDNA, barcodes ligated to transcripts/targets. | Imaging of individual pixels for proteins of interest | Extract spatially barcoded cDNA, PCR amplify with sequencing primers. | After spatial barcoding, template switch adds handles, PCR completes adapters. | Liu et al. (2020) *Cell 1* 83 (6), 1665-1681.E18. |
| **Slide-tags_recon** | Tissue sectioned | Fresh-frozen tissue section mounted on bead array; nuclei remain in situ. | na | Spatial barcodes absorbed by nuclei in situ. Spatially tagged nuclei isolated; microfluidics capture and barcoding of individual nuclei. | na | na | Downstream single-cell library amplification (standard droplet scRNA or other). | Standard Illumina adapter addition during library prep. | Russel et al. (2024) *Nature* 625, 101-109. |
| **Imaging-based spatial transcriptomic assays** | | | | | | | | | |
| | TI → SE | SE → DC | DC | DC → BC | | | | | |
| **MERFISH** | Tissue sectioned | Fixed tissue mounted on slides | Optional nuclear staining, anatomic staining and imaging | Combinatorial optical RNA barcodes: sequential rounds of hybridization and imaging | | | na | na | Chen et al. (2015) *Science* 348, aaa6090. DOI:10.1126/science.aaa6090 |

**Table S2. Mapping assay-specific methods to generalized experimental workflow steps.** Assay-specific methods were mapped to a shared set of generalized experimental workflow steps (as shown in Fig. 2) to support standardized metadata representation across single-cell, multi-omic and spatial transcriptomics data types. Each row corresponds to an assay family, and each column represents a generalized process category in the metadata model.
